# A Maxent modelling with a geospatial approach for the Habitat suitability of Flamingos in an Evanescing Ramsar site (Sambhar Lake, India) over the changing climatic scenarios

**DOI:** 10.1101/737056

**Authors:** Laxmikant Sharma Kshitij Divyansh, Alok Raj

## Abstract

Wetlands play a crucial role in the biosphere and provide numerous services. They performed multiple functions such as groundwater recharge, water purification, conservation of biological resources, act as a carbon sink and habitat of amphibians and birds. A Ramsar site-the Sambhar Lake is one of the largest inland saline wetland present in the arid region of Rajasthan, India has unique habitat suitability for the winter avifauna migrants like flamingoes and falcons. The occurrence of suitable climatic conditions and food availability like brine shrimps (*Artemia salina*) attracts flocks of migratory birds. From the last three decades, Sambhar Lake has been continuously facing degradation due to anthropogenic activities, which disturb Lake’s natural ecology and existence. These cause disturbances in habitat suitability of migratory birds in the Sambhar Lake, which leads to a reduction of population density of migratory birds. Therefore, this study aimed to assess the degradation and vulnerability of Sambhar Lake and the habitat suitability of migratory birds using Maxent Habitat Suitability model. This model provides a platform to integrate the bird’s occurrence data with the bioclimatic variables using remote sensing and Geographical Information System, and provides bird’s habitat suitability as well as predicts future bird’s occurrence scenarios. Landsat-5 and Sentinel-2 imagery for the year 1996 and 2019 respectively were used in this study. Four indicators such as LULC NDWI, MSI and SABI depicts the environmental condition of the Sambhar Lake. Output form Maxent model reveals that the Sambhar lake area with increasing anthropogenic activities has become unsuitable for flamingos. A remarkable loss of breeding sites of animal, particularly avian fauna (flamingos) is seen in the recent years due to different types of threats posed on the Ramsar site. Increase in Salt crust and Vegetation area from 36.8055 to 123.837 Sq. Km. and 26.5347 to 36.857 Sq. Km. respectively have taken place. While a decrease in saline water area from 88.8309 to 19.3256 Sq. Km has been observed, within the vicinity of Sambhar Lake as clearly shown through LULC map. The future prediction of the distribution of species in the region for the year 2050 shows that the most suitable regions will be near to *Jhapok* and nearby waters of Salt Lake City as the drains from the city opens in the lake where the flamingoes get Algae in the form of food. Active steps are needed for the lake conservation to reduce the risks of migratory bird’s population.

## 1. Introduction

Wetlands play an active role in the ecosystem and provide numerous services to humankind. They regulate the biogeochemical cycle and habitat of many amphibians and chordates. Wetlands are the areas which include (water + land) and are highly ecologically robust and productive nature. Since 1900, an estimated 64% of the world’s wetlands have already disappeared. Asia attracts major groups of migratory waterbird species, which includes threaten and least concerned birds species. Therefore, mainly migratory birds attract in after monsoon period in Asia continent; where, have had stay mainly in south Asian country like as India, Bangladesh, Pakistan, Indonesia, and Myanmar (Childress et al., 2008). The wetlands in an Asian subcontinent are evanescing at a higher pace, however; the overall trend of wetlands evanescing in inland type are more prone to disappearance as compared to the coastal zones in the vicinity (Ramsar.org/factsheet). In India, millions of migratory bird visit annually and reside in Rann of Kutch and different wetlands. The species of *Phoenicopterus* family are dominant avifauna visiting India in August-September and resides till winter and returns in month of March-April. The flamingoes have been listed in the least concern species by IUCN (International Union of Conservation of Nature). The census record of Sea world the lesser flamingoes numbers near 1.5 to 2.5 million over the globe and Chilean flamingo’s census counted less than 0.2 million (Johnson, 1997). Greater flamingos (*Phoenicopterus roseus*) are partially migratory as they resides as well as breed in this region and are distributed in the brackish saline water bodies (A. R. Johnson, 1997; Kahl, 1975) while lesser flamingos (*Phoenicopterus minor*) are affable (Brown & Britton, 1980), long-lived birds which breeds in colonies and are dispersed all over the salt pans. Greater and lesser flamingos can be differentiated easily, as the greater flamingo having black-tipped grey beak, whitish eyes and white body colour whereas the lesser flamingos are smaller in size having red eyes and dark beak. The height of Lesser Flamingos is around 80 - 90 cm, while the height of Greater Flamingo is around 110-150 cm (Sea World, 2016,). These birds usually feed in the water of 5 to 50 cm in depth (Nita et al., 2015). Dependability on the wetlands for the sustenance of the species in the arid region of India is higher as compared to the other region, due to high evaporation and lower precipitation rates. The flocks of migratory birds come in a wetland of Rajasthan. The largest inland saline wetland in India, Sambhar Lake is known as a salt lake because of its salinity is higher than 5-g/L. Here migratory birds have been coming in the huge count for past several decades; therefore, it was declared as the Ramsar site in 1990 due to its geomorphology and habitat favourability. The large region of the wetland supports a large population of flamingoes and some 71 unique types of the ecologically important avifauna. Sporadically the flamingoes are observed to breed in this region due to the climatic adaptability in the past some decades. Being a unique ecologically important habitat for the Avian winter migrants, due to the presence of the brine shrimp *Artemia salina* (Linnaeus, 1758), nowadays it has been facing challenges for its very existence. The lake has been evanescing due to the illegal salt production, excess groundwater extraction and grazing, since the last two decades. The anthropogenic activities in recent years such as disturbance in the catchment area, removal of topsoil from the lake bed, the establishment of private salt industries, poaching, are intensifying the adversity. This large legged wader is miraculous which has a very primitive pattern of nesting in the mud moulds over the floors of the shallow saline wetland. The vast expanse of sambhar lake and food availability provides a great opportunity for their breeding in the region. The sambhar lake, a Ramsar site has been struggling for its existence and has been evanescing since last two decades. The flocking of the lesser flamingoes in the Ramsar site shares a high percentage of the foreign avifauna visiting the site. The wetland has 20,000 or more count of migratory birds came in a wetland and have supported more than 1% water birds of specific species or subspecies, where that region considered as Ramsar Site under Ramsar convention guidelines. To refrain from Sambhar Lake being a Ramsar site of international importance, conservation of the lesser flamingoes is crucial. Projecting the likely effect of environmental change on the waterfowl species distributions is one of the approaches developed to understand better and counteract negative impacts of climate change (Thuiller & Munkemuller,). Habitat suitability modelling aims at defining, the ‘envelope’ that best describes its spatial range limits by identifying those environmental distributions for any chosen species. Environmental variables can be necessary for the species of interest. For bird species, commonly used variables are measures of climate (e.g. Temperature, precipitation), landscape structure (e.g. Connectivity indices), landscape heterogeneity (e.g. Ecotone cover), resources (e.g. Fishes, insects) and biotic information (e.g. Co-occurring competitor) (Thuiller & Munkemuller). Environmental variables exert direct or indirect effects on species of interest and there are significant influences under three classes (1) Limiting factors, defined as factors controlling the ecophysiology of the species (e.g. Minimum temperature) or appearance (e.g. Competition), (2) Disturbances, defined as all types of perturbations affecting environmental systems (e.g. Anthropogenic activities), (3) Resources, defined as all materials that can be assimilated by organisms (e.g. red algae, fishes, shrimps). Maxent is a general-purpose machine learning method with a simple and precise mathematical formulation, and its various aspects make it well-suited for species distribution modelling. The occurrence data of the lesser flamingo is used in establishing the correlation with the environmental variables. Maximum entropy model is used in this research for the habitat suitability of the lesser flamingo (*Phoeniconaias minor*) and greater flamingoes (*Phoenicopterus roseus*) with the Bioclimatic variables (1970-2000) WORLDCLIM-2.

## 2. Method and Material

### 2.1 Study Area

In this research the study site is sambhar inland saline wetland (lake) of state Rajasthan, (India) of which geographical coordinate is 26° 52’ N, 27° 02’ N and 74° 54’ E, 75° 14’E and its shape are tilted elliptical with an area 230 Sq. Km. The largest saline lake of India spread in three districts (Jaipur, Ajmer, and Nagaur). In the east side of the lake, one hillock of Aravalli Range situated which attained 700m height has spread northern and southern catchment. Its existence is archaic, then in an 18^th^ century here establish salt factory by British, that nowadays flourishes the socioeconomic condition of small Sambhar town. The lake basin of two divisions by a 5.16 Km. long dam near the settlements in *Jhapok* towards south and *Gudha* in the north. The water from the vast shimmering western part of the lake is pumped into the other side via sluice salt gates when it reaches, an optimum degree of salinity required for salt extraction. The saltpans can be approached by the narrow mud bunds that separate them. Some seasonal river channel support and maintain the water label of the Lake. Its climatic condition is arid, where low annual precipitation see in (fig.2) and mean temperature rises above 40°C in summer and winter fall to 10°C sometimes in May-June rise to 49°C in a year. This wetland declared as Ramsar Site 13^th^ March 1990 under guidelines of Ramsar convention (Envis Database, 2019), here come huge numbers of flocks of migratory birds such as flamingoes *(Phoenicopterus)*, Siberian crane Brahminy Shelduck *(Tadornaferruginea)*, Pacific Golden-Plover *(Pluvialis fulva)* migratory birds come at sambhar lake due to availability of food as Algae shrimps and its climatic condition is suitable between July to February month. The flamingoes are two types greater, and smaller flamingoes have distinct feature. This *Phoenicopterus* family migratory birds have migrated to specific climatic which have feasible for habitat, its length 90 cm to 150 cm (Childress, et al. 2008).

**Fig. 1:**
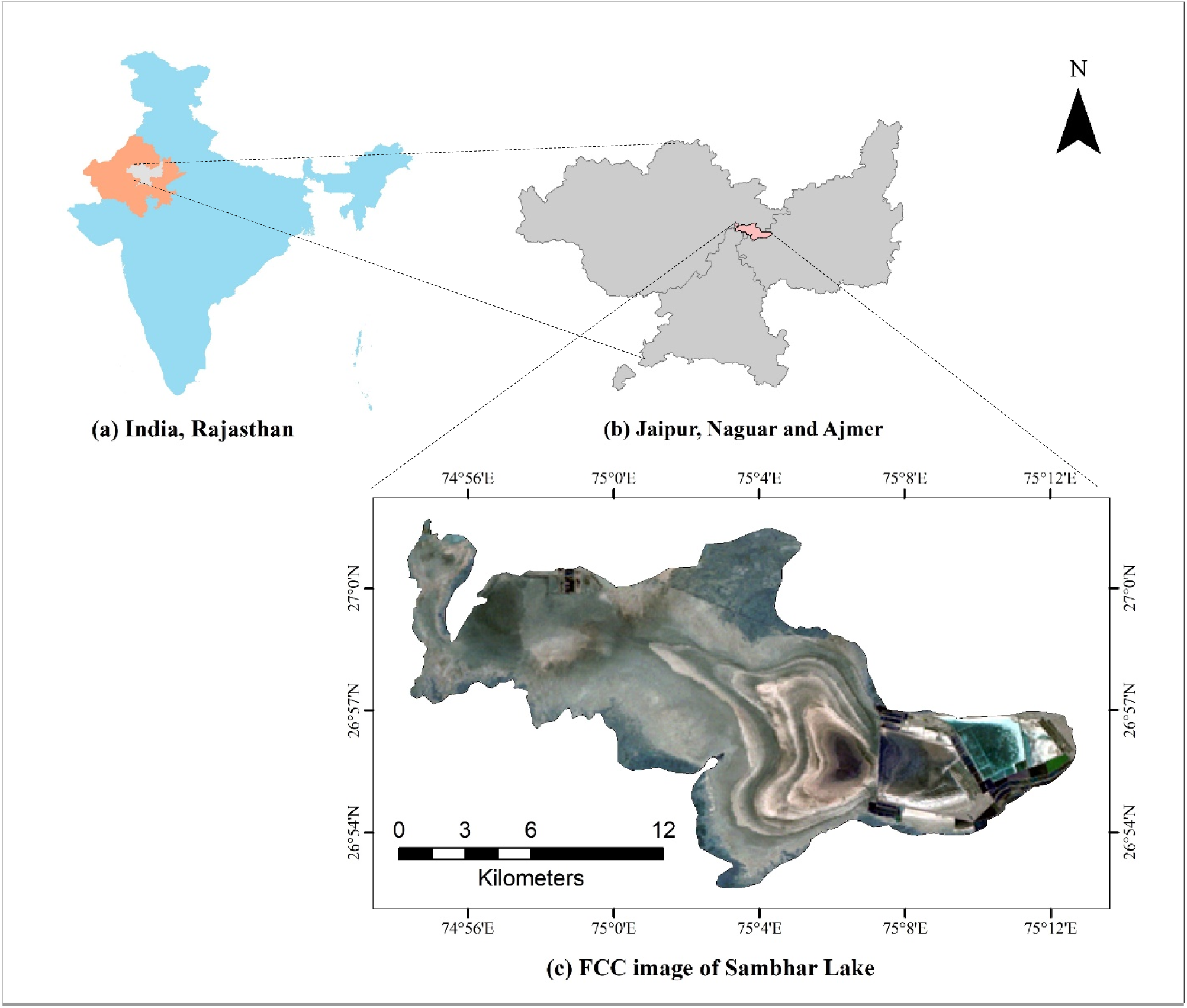
Study area map (a) Rajasthan delineating in India (b) Sambhar spreads districts Jaipur, Ajmer and Nagaur (c) FCC image of Sambhar Lake.

**Fig. 2:**
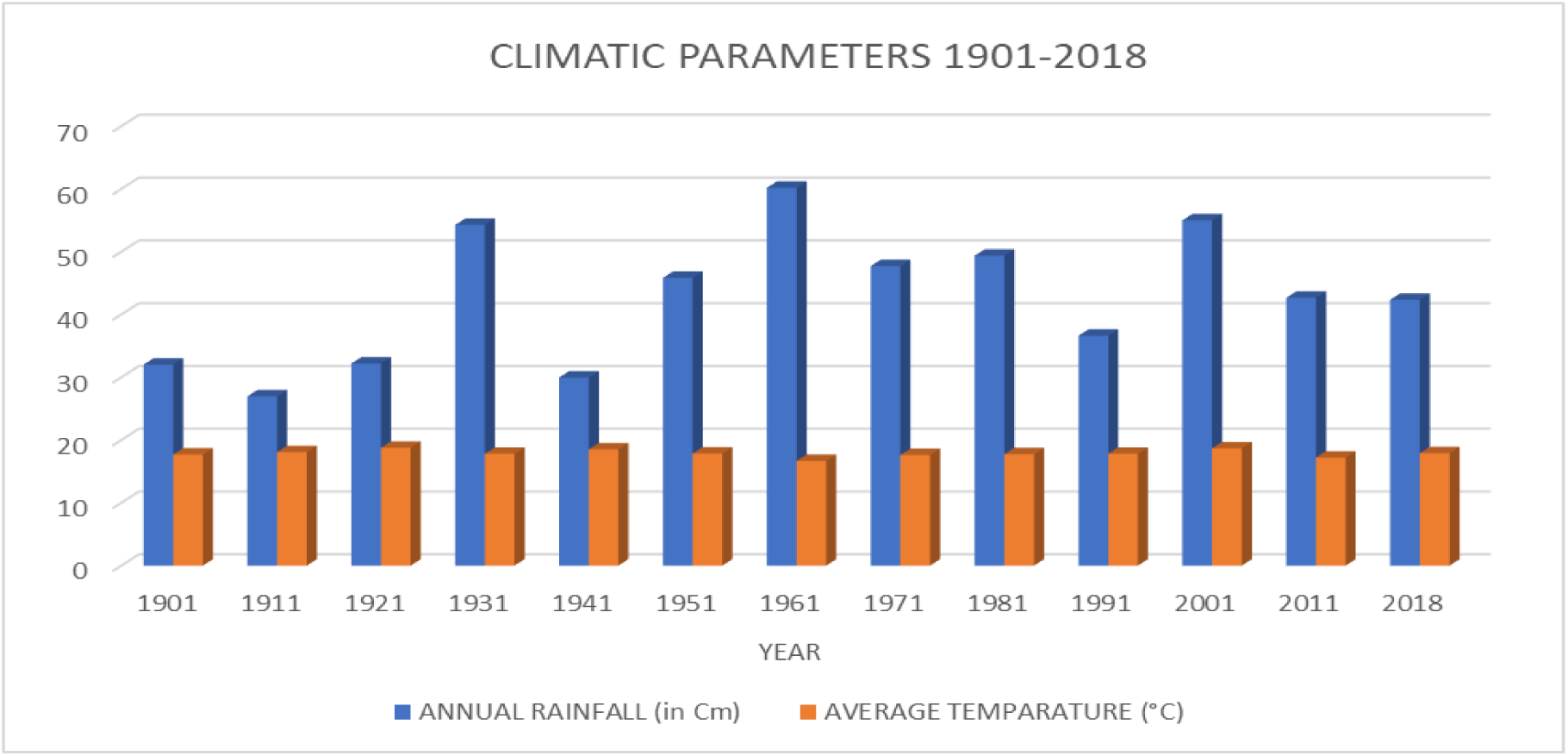
Graph illustrates the rainfall and temperature of 1901 to 2018.

### 2.2 Materials and Methodology

For assessment of the Sambhar Lake using a geospatial approach for decadal assessment and migratory bird habitat modelling using Maxent habitat model for projection of bird distribution. For assessment of lake preprocessing the satellite imagery using geospatial software like as ArcGIS 10.5 and Imagine ERDAS 15.0. Also, for creation of AOI used toposheet of 1956 of scale 1:250,000. Then the AOI used for spectral Indices (MSI, SABI and NDWI) generation and classification process through Mahalanobis distance classification) in different classes their accuracy assessment on based on ground control point of field data. For the modelling part used Maxent model; had been run based on different parameters like as bioclimatic variables, bird’s census data and spatial data, then it will give output. Based on assessment and modelling avowed the consequences over Sambhar lake and coming migratory birds.

Survey of India toposheet of 1954 of scale 1:250,000 have been acquired from Sambhar Salt-Limited of the study area. Landsat - 4/5 Thematic Mapper satellite imageries were obtained from USGS earth explorer with a spatial resolution of 30m of the year 1991, and Sentinel −2A satellite imageries were also obtained from USGS earth explorer with a spatial resolution of 10m of the year 2019. List of input data is provided in table no.1

**Table 1:**
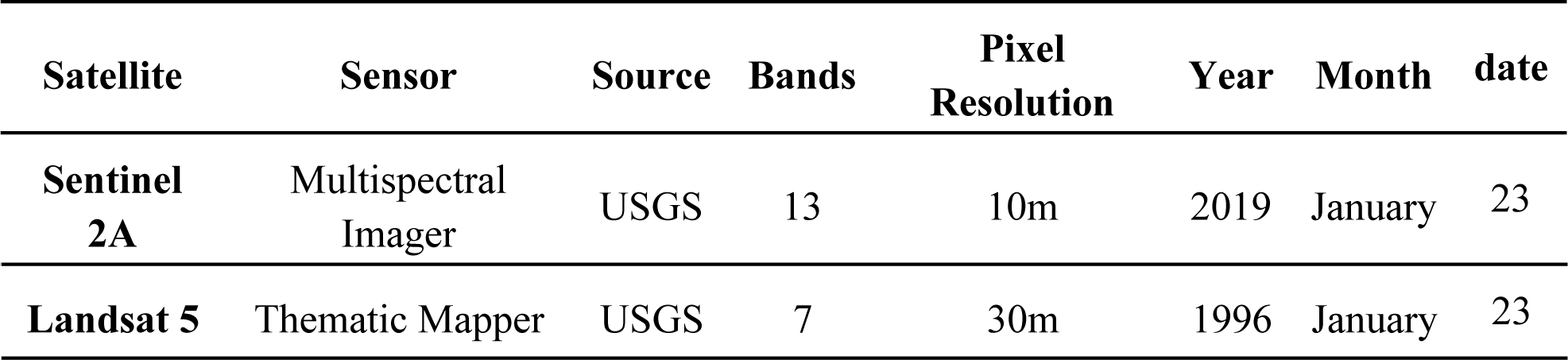
Satellite data used for LULC Classification.

The toposheet were geo-referenced based on the coordinate information given in the maps. Using ArcMap 10.5, ArcView 10.5 and Imagine ERDAS 15.0 the thematic layers (catchment area, roads, railway network, water bodies, and other) were digitised from the Topo-sheets to create a spatial database for the area of interest. The satellite imageries were geometrically corrected and referenced with the help of the toposheet by process of the map to image geo-rectification. The rectified images were mosaic, an area of interest was clipped subsequently from the mosaiced image based on the shapefile using Clip tool in Arc Map 10.5.

### 2.3 Land Use/Land Cover Classification

Land Use Land cover classification is a technique which provides the feature information through remotely sensed imagery, which has including classes like; water bodies, vegetation, bare soil and anthropogenic structures. The satellite data was processed to obtain the LULC classification for the study area for 1996 and 2019 based on visual interpretation technique and field survey using Arc Map 10.5. The mosaic images were first classified using a supervised classification technique of Mahalanobis classification, and then the classified output was verified using the respective satellite imageries. Settlements, water bodies, salt pans were classified using the method, and the changes in the land use were recorded in the subsequent years. The LULC was verified in the field, the survey of the study area was conducted in the region, and ground-truthing was done using GPS. The field survey was done in the winter season, the most favourable season for the arrival and nesting of the flamingoes. Random points were selected, and at each location, the land use pattern and coordinate information were recorded using the global position system (GPS) and the information was used for the accuracy assessment. The information collected was incorporated in the GIS domain and overlaid on the LULC map for accuracy.

### 2.4 Spectral Indices

The spectral indices are scientific, mathematical amalgamation for the satellite imagery that provides adequate information about the earth feature from the imagery. It is also indicators of environmental change. Therefore, using three different indices (NDWI, SABI, MSI)

#### Normalised Difference Water Index (NDWI)

The index NDWI is water content analysis through the combination of satellite band imageries. The band combination of NIR and Green that shows the reflectance of saline water bodies. In most cases, the NDWI can enhance the water information effectively. It is sensitive to built-up land and often results in over-estimated water bodies, but in the case of AOI being a water body only; it does not deviate with the results.

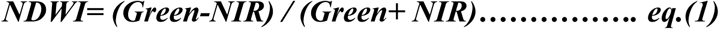

The reflectance of water in NDWI is maximised by using green band wavelengths and minimised by low reflectance of NIR by absorbing maximum of wavelength. The water features are enhanced owing to having positive values and vegetation and soil are suppressed due to having zeroed or negative values resultantly.

#### Moisture Stress Index (MSI)

It is the simple spectral ratio defining the qualitative status of the moisture content of a region. The spectral ratio assigned in an algorithm uses the two bands of the electromagnetic Waves, i.e. SWIRI and NIR. The MSI values range from 0.3 to 2.0, where the values nearing to 0.3 shows the lesser stressed region and the highly stressed regions are depicted by the values near to 2.

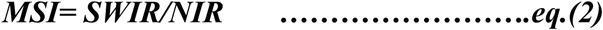

For Landsat 4/5 TM, the band’s images were selected for SWIR1-Band 5 of wavelength (1.55 nm - 1.75 nm) and NIR – Band 4 of wavelength (0.76 nm - 0.90 nm).

#### Surface Algal Bloom Index (SABI)

It is an algorithm developed for the detection of the water floating biomass which has a similar response in NIR to that of land vegetation, with some specific inclusions of water-sensitive spectral bands, Blue being characteristic of clear water and green being the one for water column blooms.

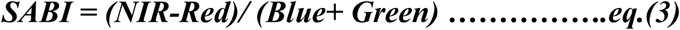

For Landsat - 4/5 TM, band images were selected for NIR-Band 4 of wavelength (0.76 - 0.90 nm), Red-Band 3 of wavelength (1.55 - 1.75 nm), Blue-Band 1 of wavelength (0.45 - 0.52 nm), Green-Band 2 of wavelength (0.52 - 0.60 nm) and for Sentinel 2A, band images were selected for NIR-Band 8 of wavelength (0.84 nm), Red-Band 4 of wavelength (0.65 nm), Blue-Band 2 of wavelength (0.49 nm), Green-Band 3 of wavelength (0.560 µm).

### 2.4 Habitat Suitability Modelling

Numerous environmental conditions are used for the determination of the habitat suitability of any target species in any region. The Habitat suitability modelling is the method of understanding the correlation between the various environmental variables and occurrence of the species as a demarcation of their most suitable habitat. For determination of which set of the environmental variables are the best fit for the habitat suitability prediction of the target species or the favourable condition under which species can sustain life. In this study, the habitat suitability is predicted for both the types of flamingoes which visit the Sambhar Lake, i.e. Greater Flamingo and Lesser Flamingo. The presence or occurrence of the flamingo is determined by the presence of saline water/ salt brine along with the other environmental and climatic conditions, which are key factors for their survival and sustenance. Maxent is a general-purpose machine learning method with a simple and precise mathematical formulation, and its various aspects make it well-suited for species distribution modelling. Predictive modelling for the spatial distribution of the species based on the environmental conditions of the given sites and the known occurrence points of the species is an efficient technique which is generally used in analytical biology, conservational reserve planning, ecological and evolutional trait and trend analysis, epidemiology, invasive-species management and various other fields. In the Maxent, occurrence data (presence only) of the target species in the region of interest is collected through various sources, i.e. GBIF, AWC census, NHBF. In the latitude-longitude format, this is tabulated in the MS Excel and saved in the comma-delimited CSV (.csv) format for the input in the Maxent. The environmental variables (e.g. temperature, precipitation) are given as input across a user-defined area of interest divided into grid cells. Maxent extracts a sample of background location (unknown presence) that it contrasts against the presence locations. The model provides a highly non-linear response curve. The bioclimatic variables given in the input was download from World-Clim (Version 2.0). Through the Habitat suitability Model (Maxent) used for future prediction of flamingoes in the sambhar lake, while using various type of data such as occurrence date/ presence data of species and future bioclimatic variable of IPCC5 of global climate model was used for the representative pathway of 4.5 for the year 2050 and 2070 climate projection. The RCP-4.5 assumes that the GHG emissions will peak till the year 2040, and then the emissions will decline further.

## 3. Results and Discussion

The LULC of the satellite image of January 1996 showed in the (Fig.5), and LULC map of January 2019 is shown in (fig.4), five classes were taken into consideration, namely-Saline Water, Salty Brine, Vegetation, Salt-pans and salt crust. The LULC was carried out, taking five classes of the radiometrically corrected satellite image and the statistical data of each class shown in the (table. 2). The supervised classification was done by providing a signature file for both the satellite images and Mahalanobis classification was done, and LULC class was analysed. The change results observed from the year 1996 - 2019 of saline water class was decreased 88.83 Sq. Km. to 19.32 Sq. Km. Therefore, saline water decreased by 69 Sq. Km. The salty brine class has decreased from 26.47 Sq. Km. to 2.29 Sq. Km. the area between 1996 - 2019, here shrinkage of brine salt area by 24 Sq. Km. Vegetation class has increased from 26.53 to 36.85 Sq. Km. between 1996-2019, observed change is 10.32 Sq. Km. In pan salt, class decreased from 45.70 Sq. Km. to 42.04 Sq. Km. and observed change is 3.66 Sq. Km. The significant change in salt crust class has observed during 1996 - 2019 from 123.83 to 36.80 Sq. Km is decreased by 87.03 Sq. Km.

**Fig. 3:**
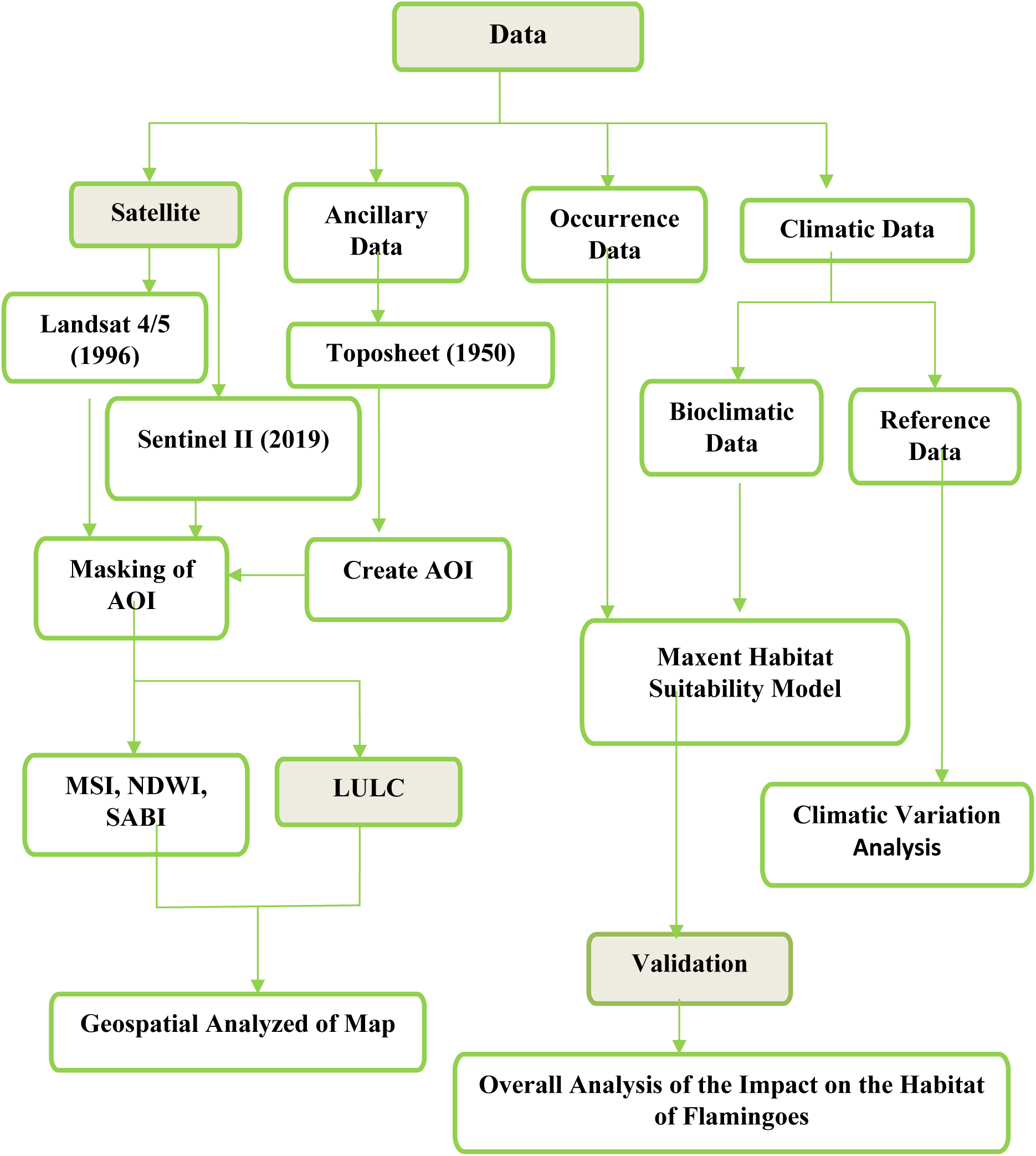
Paradigm of the overall methodology.

**Fig. 4:**
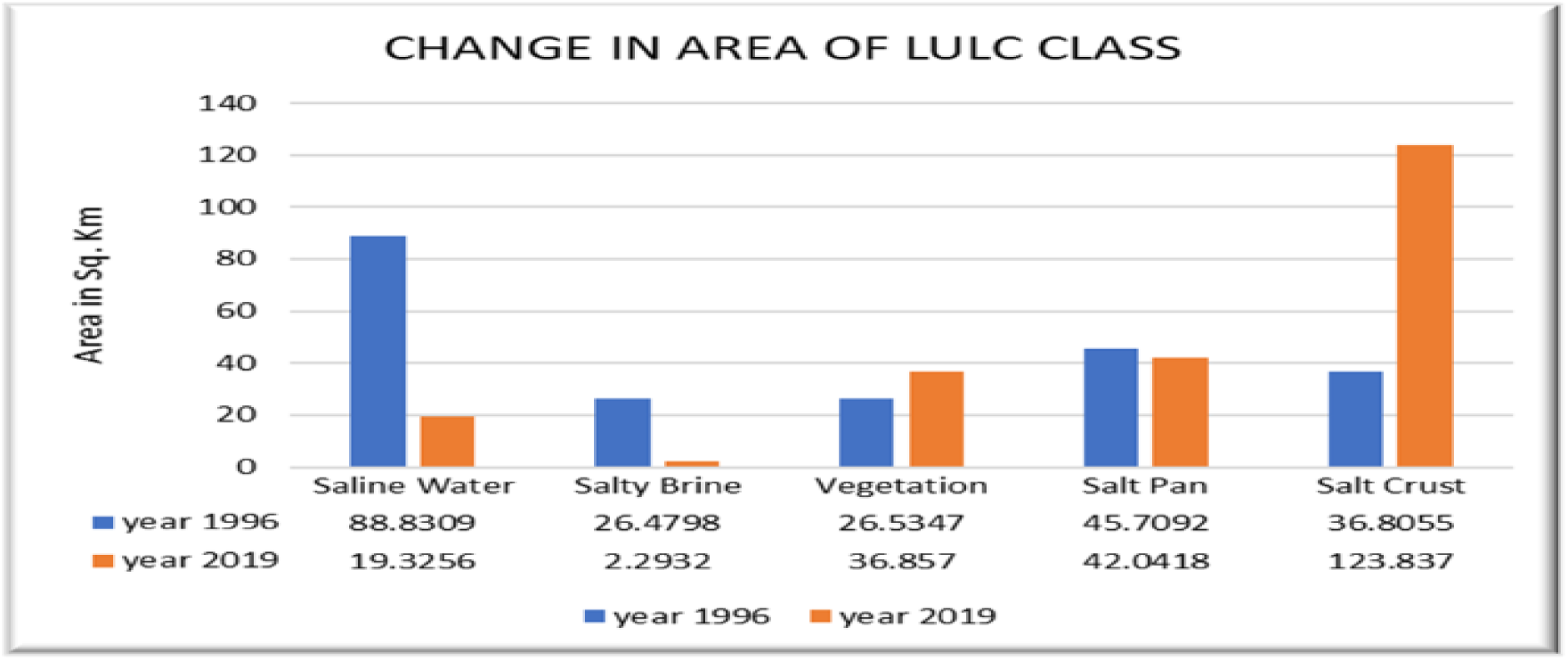
LULC area change in Sambhar Lake in 1996 and 2019.

**Fig. 5:**
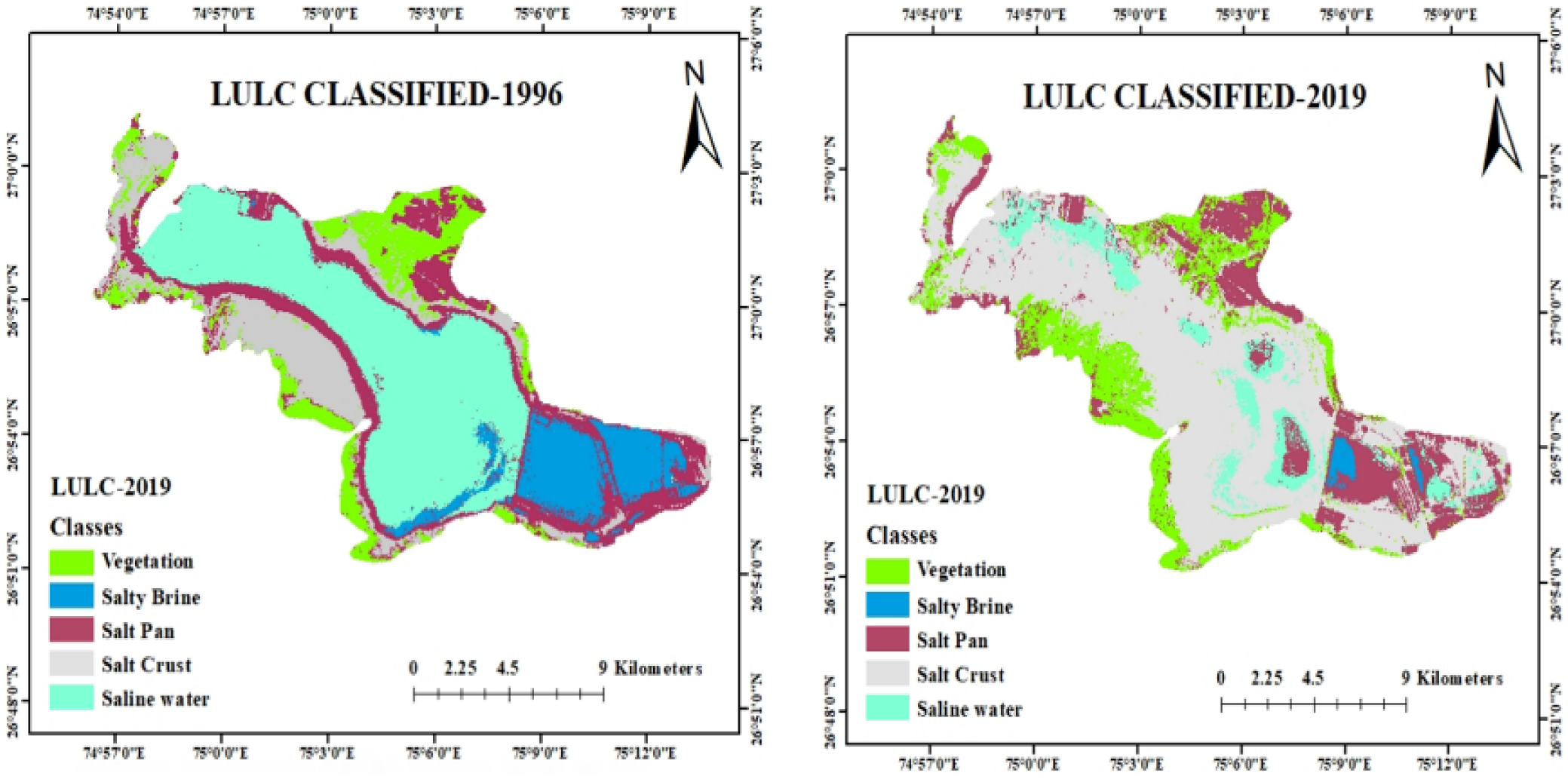
LULC of Study area for the year 1996 and 2019.

**Table 2:** Area Calculation and change for the year 1996 & 2019.

| LULC Classes | Area (Sq. Km)1996 | Area (Sq. Km.)2019 | Change (Sq. Km) |
| --- | --- | --- | --- |
| Saline Water | 88.8309 | 19.3256 | -69.5053 |
| Salty Brine | 26.4798 | 2.2932 | -24.1866 |
| Vegetation | 26.5347 | 36.857 | 10.3223 |
| Salt Pan | 45.7092 | 42.0418 | -3.6674 |
| Salt Crust | 36.8055 | 123.837 | 87.0315 |

**Table 3:** Illustrate the accuracy of the classification with a kappa coefficient.

| Year of Imagery | Classification<br>Methods | Accuracy<br>Assessment | Kappa coefficient |
| --- | --- | --- | --- |
| 1996 | Mahalanobis | 85.71% | 0.8082 |
| 2019 | Mahalanobis | 84.21% | 0.8081 |

Accuracy assessment was performed to check the number of pixels labelled correctly in the classification. It is based on reference pixels. Due to incomplete atmospheric correction, there are chances of lots of errors. Ground reference data were collected to generate the error matrix. Google earth image was used to perform an accuracy assessment in ERDAS imagine software.

### 3.1 Maxent Habitat Suitability Modelling

The resulting map produced from the Maxent model for flamingo is shown in (Fig.6). In the map, warmer colours represent the predicted suitable habitat, whereas the colder colours show the least suitable habitat for flamingos. White dots represent the occurrence data used as an input while the violet dots show the test locations. Habitat suitability value per pixel (per cell) was given by the model between 0 to 1, with 0 values being not suitable while one is highly suitable. Habitat suitability map depicts that most of the predicted flamingo habitat is distributed nearby wetlands. Blue pixels are showing area not suitable (0.0091 - 0.2), cyan pixel for least suitable (0.2 - 0.4), green pixel for moderately suitable (0.4 - 0.6), orange pixel for suitable (0.6 - 0.8), and red pixel for highly suitable (0.8 - 1.0) habitat for flamingos. (fig.6) represents the occurrence of the flamingoes in the Sambhar Lake, dating back from 1885 to 2019 and the future distributions of the species are projected.

**Fig. 6:**
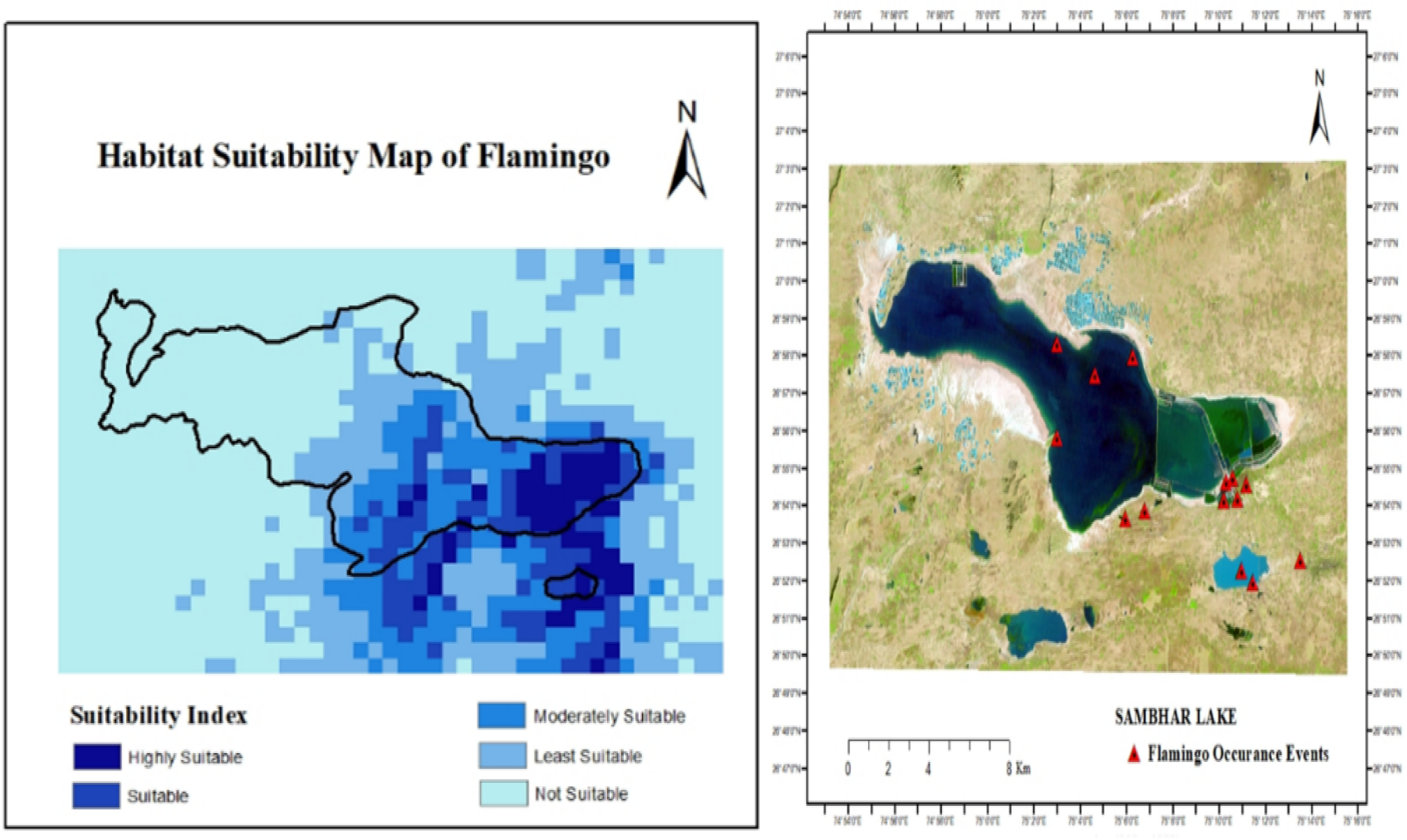
Habitat Suitability map for present condition and Occurrence Map of the flamingo in Sambhar Lake.

### 3.2 Habitat Suitability of the Flamingoes in Future

Fig.6 represents the future projection of the species distribution for the future years 2050 & 2070. The data used here are the IPPC5 climate projections from global climate models (GCMs) for the representative concentration pathways (RCPs) RCP-4.5. The data at 30-seconds (of a longitude/latitude degree) spatial resolution was obtained in the GeoTIFF file format, for the period 2050, the average is taken for 2041 - 2060 and period 2070, the average is taken for 2061 - 2080. These are the most recent GCM climate projections that are used in the Fifth Assessment IPCC report. The GCM output was downscaled and calibrated (bias-corrected) using WorldClim-1.4 as baseline ‘current’ climate. Habitat suitability map (Fig.6) depicts that most of the predicted flamingo habitat is distributed nearby the wetlands. Light Blue pixels are showing area not suitable (0.0091 - 0.2), Carolina blue pixel for least suitable (0.2 - 0.4), Azure blue pixel for moderately suitable (0.4 - 0.6), Navy blue pixel for suitable (0.6 - 0.8), and Royal blue pixel for highly suitable (0.8 - 1) habitat for flamingos. The future projection for the distribution of the Flamingo species in the Sambhar lake for the year 2050 having CMIP-5 future GCM. The highly suitable region in the year 2050 has reduced to a greater extent as compared to the recent year habitat suitability. In the year 2050, the RCP-4.5 has been used, and it is assumed that the Greenhouse gas emission will be at peak in the year 2040 and then the emission will reduce; this may lead to the rise in temperature in the region and decrement in the rate of precipitation. Has been reducing which

The environmental conditions in the region may become adverse if the climate changes at this pace. The habitat of flamingoes will become unsuitable and lead to migration of flamingo species from the region, which can also affect the number of various migratory avifaunas which may not sustain life in the adverse environmental conditions. The average temperature of the region has increased by 1 - 2 °C from the year 1901 to 2018. The rate of precipitation (seasonality) and annual rainfall in the region have decreased in a century. Habitat suitability map (fig.7) depicts that most of the predicted flamingo habitat is distributed nearby the wetlands. Light Blue pixels are showing area not suitable (0.0091 - 0.2), Carolina blue pixel for least suitable (0.2 - 0.4), Azure blue pixel for moderately suitable (0.4 - 0.6), Navy blue pixel for suitable (0.6 - 0.8), and Royal blue pixel for highly suitable (0.8 - 1) habitat for flamingos. The future projection for the distribution of the Flamingo species in the Sambhar lake for the year 2070 having CMIP-5 future GCM. The highly suitable region in the year 2070 has reduced to a greater extent as compared to the recent year habitat suitability. In the year 2070, the RCP-4.5 has been used, and it is assumed that the Greenhouse gas emission will be at peak in the year 2040 and then the emission will reduce; this may lead to the rise in temperature in the region and decrement in the rate of precipitation. The Combination and indices are the dimensionless quantity that defines the qualitative significance of the feature-based upon understanding. Water indices are rationing bands, which sensed ground reflectance values of the vegetative area and water bodies. Several waters and algal presence indices in the remote sensing field have been used.

**Fig. 7:**
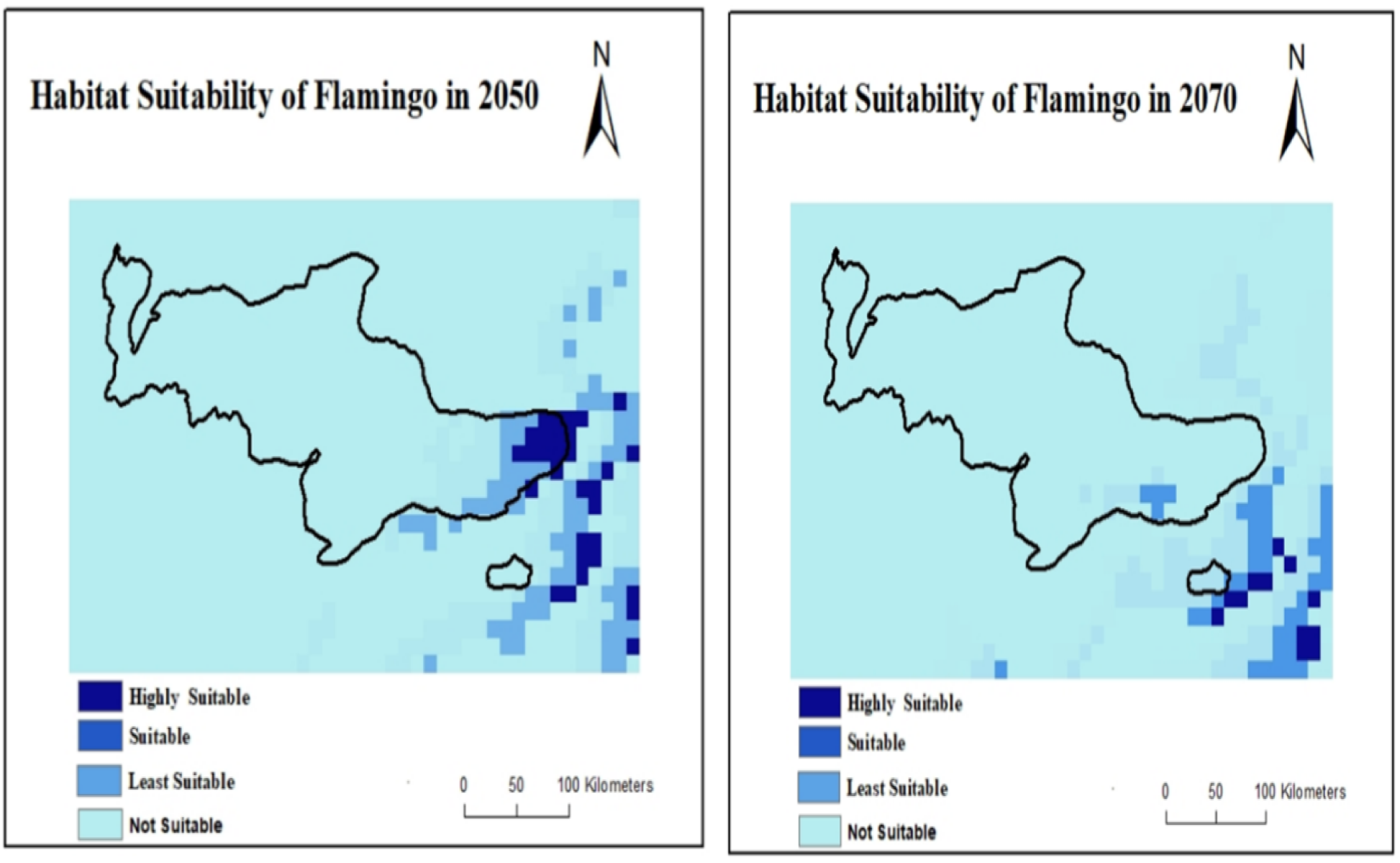
Illustrate the projection of suitable habitat of Flamingo in 2050 and 2070 Geospatial Analysis of the Sambhar Lake.

Various indices used for the analysis of the condition of Sambhar Lake and the change in the past two decades. NDWI of the year 1996 ranged from 1.0 to −0.1 in which the saline water had the value near to 1.0, which represents the presence of water in the region and zero to negative values represents the land/soil and vegetation. (fig.8) Illustrates that higher water concentration in the lake due to the heavy rainfall in the year 1994 & 1996 in the region. Eastern region of the lake has the dam, and it is used for the salt production which has a slightly lower concentration of the water. The salt pans had the values near to 0 representing the least water concentration in the lake region due to the presence of brine in it for the salt extraction. The NDWI for the year 2019 (fig.8) ranged from 1.0 to −0.1, in which the salt pans and some other area of depression only have values near to 1 representing the presence of water. The region of Salty Brine (Dam) has values ranging near to 0.5, which represents the lesser presence of water. The water level in Sambhar Lake has drastically decreased in the past two decades, and the water index (NDWI) depicts the depletion trend. The water index MSI (Moisture Stress Index) for the year 1996 ranged from 0.1 to 3.0 in which the values near to 0.1 represents the least stressed region and values near to or greater than 2 represents the highly stressed region. The moisture in the region was higher near to the catchment, lake, dam and salt pans due to the presence of water in the area. The MSI for the year 1996 had values near to 0.1 for the significant region of catchment and dam, depicting the lower stressed condition in the region and values nearing to 3.0 showed the salt crust and land/ saline soil region having the highly stressed condition. MSI of the year 2019 (fig.9) value ranges from 0.1 to 3, where the region near the dam and salt pans are showing values near to 0.1 and having less moisture stressed condition, whereas the western region of the lake has values near to 3, depicting highly stressed condition. The Moisture stress for the Sambhar Lake region has increased abruptly in the last two decades, and the water catchment area has been decreasing.

**Fig. 8:**
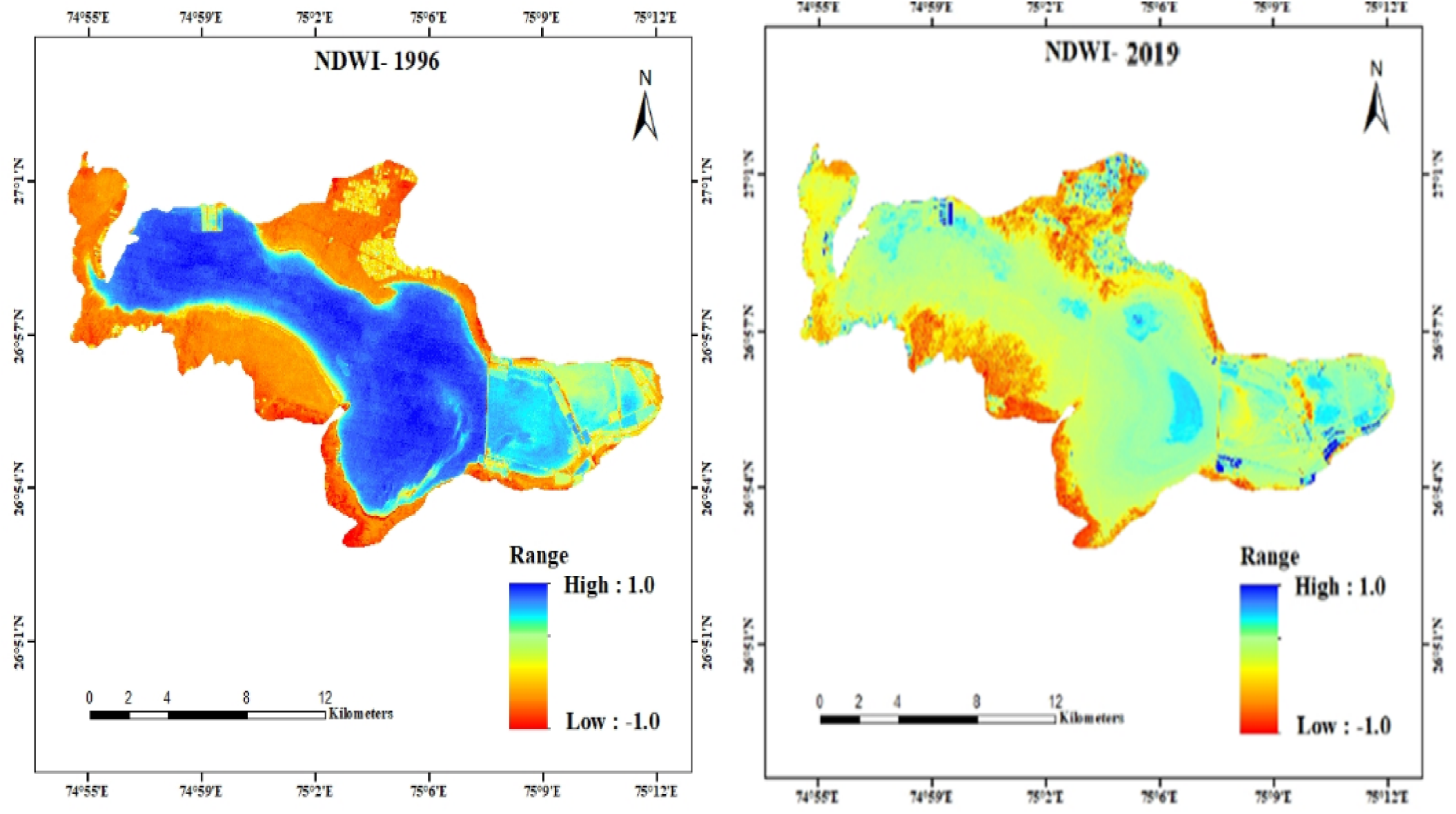
Map of Water Index NDWI (1996 and 2019)

The wetland region harms the egg-laying regions of the flamingoes and breeding is being affected, resultant of the reduction in the number of migratory avifaunaeVisiting the Ramsar site. The flamingo populations rely on the wetlands for breeding and sustenance of life and water in the wetland provides them with a suitable habitat. The Surface Algal Bloom Index (SABI) is an empirical algorithm developed to detect water floating biomass. The SABI value for the Sambhar lake region ranges between −1 to 1.0, where the values nearing to −1.0 represents the presence of the water column algal blooms, values nearing to 0 represents the surface algal blooms and values nearing to 1.0 represents the absence of algal blooms and sometimes the vegetation due to presence of chlorophyll-a. The values of SABI for the year 1996 ranged from −1.0 to 1.0, where the western region of the lake had values nearing to −1.0 and had higher water column algal bloom concentration whereas the salt pans had the values nearing to 0 as the water was shallow and depicted the surface like characters. The values of SABI for the year 2019 (fig.10) ranged from −0.1 to 1.0, where the salt pan and some region of lake had values nearing to −1.0 representing presence of water column algal blooms and the western region of the lake has values nearing to 0 or above representing lesser algal blooms in the region. The surface algal bloom SABI depicts the water column algal community and surface algal community present in the region. The region having higher algal bloom concentration becomes the suitable region for the sustenance of life as the algae are the primary food source for the flamingoes and they feed upon red algae, saline microorganisms and smaller insects.

**Figure. 9:**
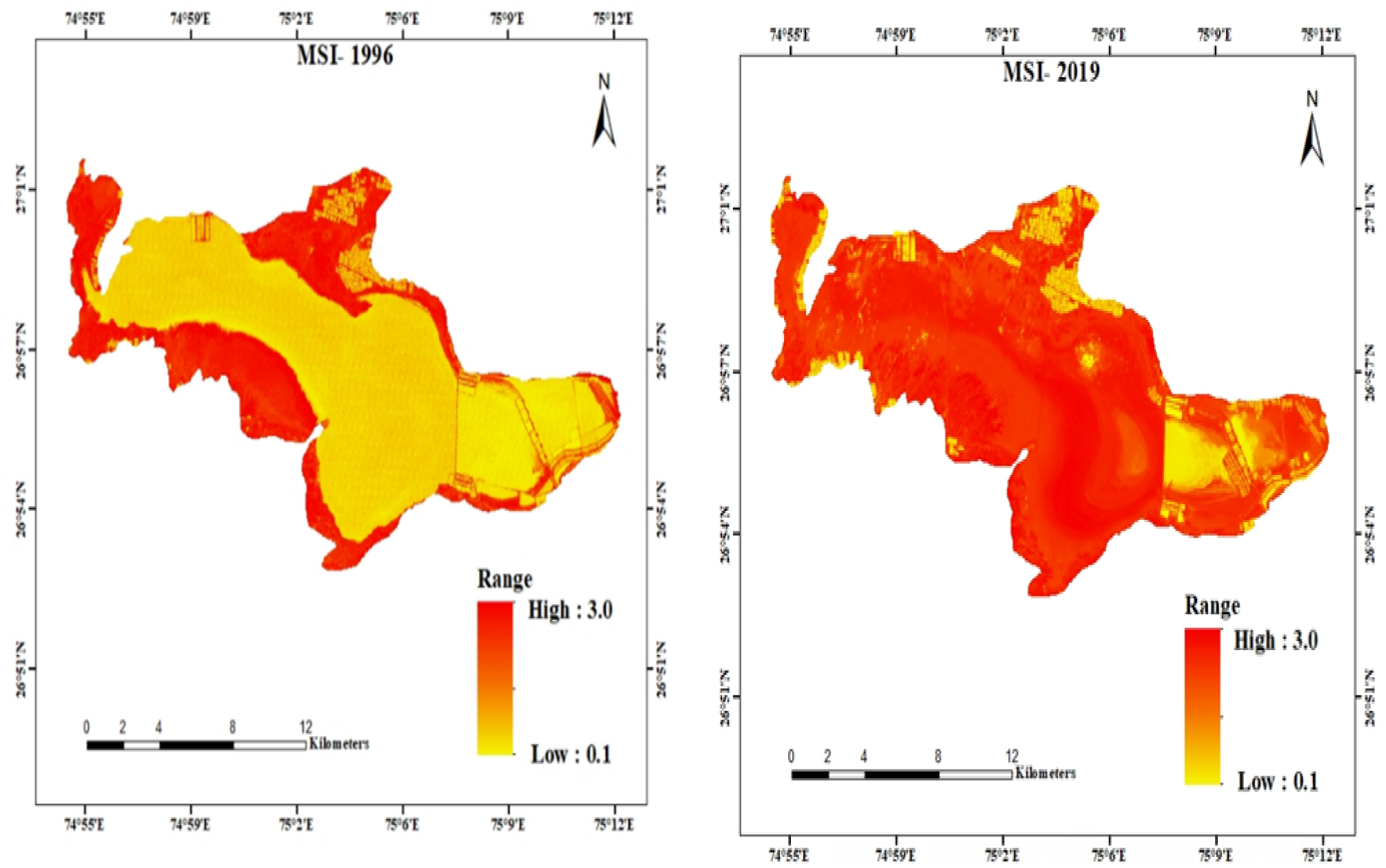
Map of moisture stress index (1996 and 2019)

**Fig. 10:**
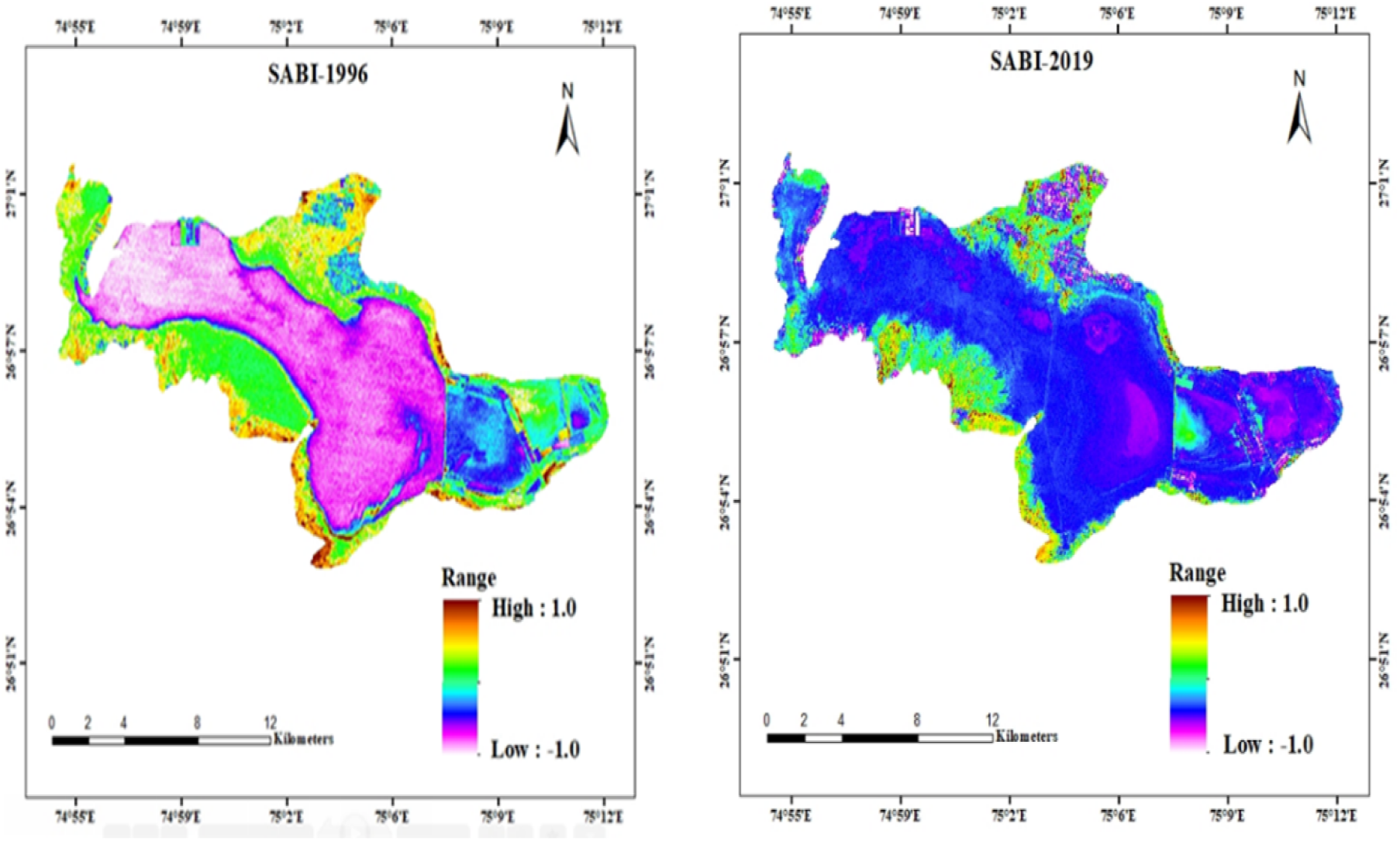
Map of Algal Bloom Index (SABI) for the year 1996 and 2019.

## 4. Conclusion

Sambhar lake has a great potential to sustain large populations of flamingoes; thus, they have high occurrence events in the past decades. LULC changes clearly show that the vegetation and salt crust have drastically increased nearby the lake. The primary reason for these changes is mushrooming of private salt industries, removal of topsoil by salt producers, construction of small dams in the catchment area and heavy vehicular trespass by villagers. A remarkable loss of breeding sites of avian fauna (flamingos) is seen in the recent years due to different types of threats posed on the Ramsar site. During period 1996 to 2019, an increase in Salt crust from 36.80 Sq. Km. to 123.83 Sq. Km. and vegetation form 26.53 Sq. Km. to 36.85 Sq. Km. respectively, while there was a decrease in saline water from 88.83 Sq. Km to 19.32 Sq. Km. From this study, it is observed that both species of flamingos (lesser and greater) stay in the lake throughout winters when the lake contains only a concentrated solution of brine. In recent years, the lake had dried up suddenly on many occasions when sluice gates were opened to drain the water into saltpans, due to which massive population of birds have had to abandon the lake in mass. This shows a drastic decrease in the population of flamingos in the last few years. Maxent (HSM) predicted that the wetland areas near the water body are most suitable for flamingos, especially areas near to the *Jhapok*, salt dam, Salt Lake City and majorly the eastern part of the lake. The higher salinity is the indicator for the presences of red algae in the lake; therefore, the occurrence of food availability is higher for flamingos, which is one of the main reasons for higher habitat suitability of flamingoes. Maxent results also show that the area with increasing anthropogenic activities has become an unsuitable habitat for flamingos. The model predicts that the most suitable regions for the distribution of flamingoes in the lake for the year 2050 will be near to *Jhapok* and drains of Salt Lake City as the drains from the city opens in the lake where the flamingoes get algae in the form of food. The projections based on bioclimatic variables show that the western side of the lake will not be suitable for the flamingoes and result in the migration of the species from the Ramsar Site and it could lose the status of Ramsar site. The future species distribution for the year 2070 depicts the clear picture of the climate change and its varying impacts on Sambhar Lake. The regions nearby the *Kochia ki Dhani* and the salt dam will be the only region being suitable and least suitable leaving the whole Sambhar Lake the non-suitable region for the breeding and sustenance of Flamingos based on the bioclimatic conditions. NDWI and MSI shows that the water concentration near the wetland has been declining in the two decades, where the water concentration was reasonably high in the lake in the year 1996 due to recent heavy rainfalls and proper recharge of the water in 1994 and the water concentration in the year 2019 was lesser. The rainfall and evapotranspiration in the region are not proportional as the higher temperature during summer quarters, due to which the availability of water is less and further extraction of groundwater and surface water for illegal salt production is worsening the scenario. The SABI shows the presence of water column algal blooms and the surface algal blooms in the Sambhar Lake. The concentration and presence of algal blooms play a determining factor for flamingo’s habitat suitability as algae is a leading food for the flamingoes. The SABI values in the year 1996 were higher and spread over a large area of the lake, but in the year 2019, this has decreased drastically to a lesser region having algal blooms due to drying up of the various regions of the lake. The trend in the reduction in the algal blooms may lead to lesser availability of food for flamingoes and lead to severe survival conditions for them.

